# Altered Gut Microbiota Mediates the Association between APOE Genotype and Amyloid-β Accumulation in Middle-Aged Adults

**DOI:** 10.1101/2025.07.27.667048

**Authors:** Yannick N. Wadop, Rebecca Bernal, Wepnyu Y. Njamnshi, Claudia L. Satizabal, Alexa Beiser, Agustin Ruiz, Alfred K. Njamnshi, Ramachandran S. Vasan, Sudha Seshadri, Jayandra Jung Himali, Bernard Fongang

## Abstract

**Importance:** The apolipoprotein E (*APOE*) ε4 allele is the strongest genetic risk factor for Alzheimer’s disease (AD), yet the mechanisms linking *APOE* to amyloid-β (Aβ) pathology remain incompletely understood. Emerging evidence suggests that the gut microbiome may modulate neurodegeneration; however, its role as a mediator in the APOE–Aβ relationship remains unclear.

**Objective:** To evaluate whether specific microbial taxa mediate APOE-related effects on brain Aβ burden in an established population-based study of middle-aged adults.

**Design, Setting, and Participants:** This cross-sectional study analyzed data from the Framingham Heart Study cohort. Data were collected at the third examination visit (n = 227, %Female = 58, mean age = 56.5 ± 8.3), between 2016 and 2019.

**Exposures:** Gut bacterial DNA was sequenced using 16S rRNA, and amplicon sequence variants (ASVs) were agglomerated at various taxonomic levels (14 phyla, 70 families, and ∼140 genera). *APOE* genotypes were derived from blood DNA using PCR and restriction isotyping. Predicted microbial functional potential was based on KEGG Orthologs.

**Main Outcomes and Measures:** Overall and regional measures of cerebral amyloid-β deposition were assessed using carbon-11 Pittsburgh (PiB) Compound-B PET scans. The global PiB deposition served as the primary outcome for “*overall*” amyloid burden. Regional amyloid deposition values were analyzed as secondary outcomes.

**Results:** A higher Aβ burden was significantly associated with the depletion of protective genera (e.g., *Faecalibacterium* β [95%CI],-0.35 [-0.40,-0.30]; *Ruminococcus*-0.25 [-0.27,-0.23]; *Butyricicoccus*-0.27 [-0.32,-0.22]) and the enrichment of pro-inflammatory taxa (e.g., *Alistipes* 0.07 [0.06, 0.08], *Bacteroides* 0.10 [0.07, 0.13]) and *Barnesiella* (0.18 [0.16, 0.20]). These associations were more pronounced in *APOE* ε4 carriers, who exhibited a broader spectrum of microbial dysbiosis. Mediation analysis showed that *Ruminococcus*, *Butyricicoccus*, *Clostridium*, and *Christensenellaceae* collectively mediated ∼0.3-0.4% of the effect of *APOE* ε4 on global Aβ burden. Functional profiling revealed a reduced abundance of microbial genes involved in key metabolic pathways among individuals with higher Aβ levels.

**Conclusion and Relevance:** Gut microbiome composition partially mediates the relationship between *APOE* ε4 and cerebral amyloid burden. These findings support a gut-brain axis mechanism in AD and suggest that microbiome-targeted interventions may mitigate *APOE*-related risk.

## Introduction

Alzheimer’s disease (AD) is neuropathologically characterized by the accumulation of extracellular amyloid-β (Aβ) plaques and intraneuronal tau neurofibrillary tangles in the brain parenchyma.^1–4^ These proteinopathies begin to accumulate years, if not decades, before the onset of clinical symptoms, as evidenced by positron emission tomography (PET) imaging in cognitively unimpaired individuals. While the pathogenic cascade of AD is multifactorial, emerging research implicates the gut microbiome as an important contributor to neurodegenerative processes.^5,6^ The gut microbiota is intimately involved in numerous aspects of human physiology, and perturbations in the microbiome have been linked to metabolic and inflammatory diseases.^7,8^ In the context of brain health, accumulating evidence suggests that shifts in gut microbiome composition are associated with neuroinflammation and the development of neurodegenerative diseases.^9^ This gut and brain connection is mediated through a bidirectional communication network known as the gut–brain axis.^10–13^ Through neural, endocrine, and immune pathways, gut microbes can influence brain physiology and pathology.^14–19^

Several cross-sectional studies and animal models support a link between altered gut microbiota and AD pathology.^20,21^ AD patients have been reported to exhibit gut microbiome profiles enriched in pro-inflammatory taxa and depleted in beneficial commensals relative to cognitively normal controls.^22,23^ Notably, taxa such as *Faecalibacterium*, *Ruminococcus*, and *Barnesiella* (typically regarded as anti-inflammatory or short-chain fatty acid producers) are reduced in AD, whereas certain pro-inflammatory genera are increased.^24^ These microbial changes correlate with elevated peripheral inflammatory markers in individuals with amyloid pathology, suggesting that the gut microbiota may modulate AD risk via immune-inflammatory mechanisms. In support of this idea, it was shown that cognitively impaired individuals with brain amyloidosis had higher abundances of pro-inflammatory gut bacteria (and lower anti-inflammatory bacteria), accompanied by increased inflammatory markers.^25^ Conversely, interventions that modulate the microbiome can impact amyloid pathology. In transgenic AD mouse models, exercise or probiotic supplementation has been shown to attenuate amyloid accumulation and reduce neuroinflammation.^26^

While the connections between the gut microbiome and AD pathology are increasingly recognized, the influence of host genetic factors on this gut-brain axis is only beginning to be explored.^27,28^ The strongest genetic risk factor for both early-onset and late-onset AD is the apolipoprotein E (*APOE*) gene polymorphism.^29,30^ The *APOE* gene has three common protein isoforms (ε2, ε3, ε4), yielding six possible genotypes in humans.^31,32^ The ε4 allele is associated with a dose-dependent increase in AD risk: carrying one ε4 allele confers an estimated ∼3-fold increased risk of developing AD (compared to the common ε3/ε3 genotype), and two ε4 alleles lead to even greater risk and earlier onset.^33–37^ In contrast, the ε2 allele is relatively protective against AD.^38^ *APOE* encodes the apolipoprotein E, a lipid-transport protein that plays a major role in lipid metabolism in both the periphery and the brain.^39,40^ *APOE* is a key carrier of cholesterol and other lipids, and in the brain, it helps clear Aβ peptides from the interstitial fluid.^38,41^ The isoform-specific effects of APOE on AD are complex; *APOE* ε*4* is thought to be less effective at Aβ clearance and to promote neuroinflammatory and synaptotoxic pathways.^38,42,43^ However, the precise mechanisms by which *APOE4* accelerates AD pathogenesis are still under intense investigation.

Importantly, host genetics can also shape the gut microbiome. Recent research indicates that *APOE* genotype is associated with specific gut microbiome profiles.^9^ For example, one study reported that humans carrying the *APOE* ε*4* allele (and *APOE*-targeted replacement mice) exhibited differences in gut microbial communities, including changes in *Lachnospiraceae*, *Ruminococcaceae*, and several SCFA-producing genera.^9^ These findings suggest that APOE4 may predispose individuals to a pro-AD gut microbiome composition (low anti-inflammatory microbes, high pro-inflammatory microbes), which could, in turn, exacerbate amyloid accumulation and neurodegeneration. This three-way interplay between host genotype, gut microbiota, and amyloid pathology has not been directly examined in humans to date.

Here, we address this gap by leveraging data from the Framingham Heart Study (FHS) to investigate whether the gut microbiome mediates the relationship between *APOE* genotype and brain Aβ burden. We hypothesized that *APOE* ε*4 carriers would exhibit a gut microbiome signature associated with higher amyloid burden*, particularly characterized by a reduction in anti-inflammatory bacteria. We performed 16S rRNA gene sequencing on stool samples from 227 cognitively normal to mildly impaired adults who also underwent Aβ PET imaging to explore the role of the gut microbiome as a potential modulator of genetic risk for AD.

## Methods

### Study Design and Participants

This study utilized data from the Framingham Heart Study (FHS), a multi-generational longitudinal cohort initiated in 1948, to investigate risk factors for cardiovascular disease.^44,45^ In recent years, the FHS has expanded to include studies on brain aging.^44,46,47^ Starting in 2015, members of the FHS Third Generation cohort were invited to participate in an amyloid imaging sub-study. Key eligibility criteria for the PET sub-study included: **(1)** age ≥ 30 years at the time of the third examination visit (2016–2019) and (**2**) absence of major neurological conditions (including clinical dementia, stroke, or multiple sclerosis). A total of 227 participants provided stool samples and underwent Aβ PET scanning approximately around the time of their third examination (2016-2019). All participants gave written informed consent, and the study protocol was approved by the Institutional Review Boards of Boston University Medical Center and Massachusetts General Hospital.

### Gut Microbiome Collection and Analysis

Participants provided a single stool sample, which was collected and processed following standardized FHS protocols.^44,48,49^ Stool specimens were immediately frozen at-80°C after collection and stored until batch analysis. Microbial DNA was extracted using the Qiagen PowerSoil DNA Isolation Kit (Qiagen, Hilden, Germany) according to the manufacturer’s guidelines. The V4 hypervariable region of the 16S rRNA gene was amplified by polymerase chain reaction and sequenced on the Illumina MiSeq platform to generate paired-end reads (2×250 bp). Quality control and amplicon sequence variant (ASV) inference were performed using the DADA2 pipeline^48^, yielding high-resolution bacterial sequence features. The taxonomic assignment of each ASV was performed against the SILVA 16S rRNA database (version 132). For downstream analyses, we agglomerated ASVs at various taxonomic levels (phylum, family, genus) and focused primarily on genus-level relative abundances. Rare taxa with extremely low counts were filtered out to reduce noise. We calculated α-diversity (within-sample diversity) metrics, such as the Shannon index, and β-diversity (between-sample differences) using Bray–Curtis distances. Although our primary analyses focused on specific taxa abundances rather than global diversity measures, we also employed these metrics.

### A**β** PET Imaging and Quantification

Participants underwent carbon-11 Pittsburgh Compound-B PET scans to measure cerebral amyloid-β deposition, performed at the Massachusetts General Hospital PET facility. Imaging was conducted on HR+ or GE Discovery MI PET scanners using standardized protocols as previously described.^50–53^ In brief, each participant received a 10-15 mCi bolus injection of [^11C]PiB, followed by dynamic PET acquisition over 60 minutes (four 5-minute frames, 50–70 min post-injection, were used for analysis). T1-weighted volumetric brain MRI scans (acquired within a year of the PET scan) were used for co-registration and definition of regions of interest. PET images were motion-corrected, spatially co-registered to each individual’s MRI, and normalized to a common space. A cerebellar gray matter reference region was used to calculate standardized uptake value ratios (SUVR) for cortical target regions.^50,54,55^ In addition, a global Aβ composite measure was computed by averaging PiB SUVR across a set of frontal, parietal, temporal, and retrosplenial cortical regions (often termed the FLR composite).^55^ This global PiB SUVR served as our primary outcome for “*overall*” amyloid burden.^55^ This global PiB SUVR served as our primary outcome for “*overall*” amyloid burden. Regional SUVR values were analyzed as secondary outcomes to explore region-specific relationships. Twelve regions were examined based on their known vulnerability to early amyloid accumulation and their susceptibility to early AD pathology (including the precuneus, inferior temporal cortex, frontal pole, accumbens area, amygdala, cingulate, insula cortex, hippocampus, entorhinal cortex, parahippocampal cortex, temporal pole, cerebellum white matter, and cerebral white matter).^55–57^ A global PiB SUVR of 1.10 is commonly considered an abnormal threshold for amyloid positivity; in our sample, the median global SUVR was around 1.06, with ∼23% of participants exceeding 1.10 (reflecting a mix of amyloid-positive and-negative individuals, as expected for this age range).

### APOE Genotyping

DNA was isolated from whole blood samples collected at FHS exams using a standard salting-out procedure.^58^ *APOE* genotyping was performed as described by Lahoz *et al*.^59^ The region of the *APOE* gene encompassing the two polymorphic sites (codons 112 and 158) was amplified by PCR.^60^ The PCR product was then digested with the restriction enzyme HhaI, which yields characteristic fragment patterns for the ε2, ε3, and ε4 alleles. Fragment separation was done on an 8% polyacrylamide gel and visualized under UV light after ethidium bromide staining. From the band patterns, we determined each participant’s APOE genotype (ε2/ε2, ε2/ε3, ε3/ε3, etc.).

For quality control, known *APOE* control samples were run in parallel, and a random 5% subset of samples was re-genotyped with 100% concordance.

### Covariates

Standard questionnaires were used to obtain demographic data at the third examination between 2016 and 2019. FHS participants reported their age and sex. The staff collected the dates at which participants provided stool samples and the dates for PET scans. A time difference between stool collection and PET scans (in weeks) was derived from both dates. Additionally, standardized protocols were used for clinical measures. Notably, height and weight for body mass index assessment (BMI; kilograms divided by height). **Statistical Analysis**

All statistical analyses were conducted using R (v4.2.3).^61^ Our overarching analytic approach was to identify associations between gut microbial features and Aβ burden, assess how these associations differ by APOE genotype, and test for mediation by microbial features in the APOE-Aβ relationship. Based on prior literature and directed acyclic graph considerations, we included the following covariates in all models: age, age squared, sex, BMI, PET scanner type (HR+ vs. Discovery), and the time interval (in weeks) between stool collection and the PET scan. These factors were chosen to account for potential confounding of microbiome-Aβ relationships (e.g., age and sex influence both gut microbiome and amyloid burden; BMI relates to systemic metabolism; different PET cameras have different signal characteristics; and the stool-to-PET time interval could introduce measurement noise).

### Association of Gut Microbiome with A**β**

To identify specific bacteria associated with amyloid burden, we used the MaAsLin2 (Multivariate Association with Linear Models) R package^62^ to perform generalized linear models for each taxon. MaAsLin2 was run at the genus level (and repeated for phylum and family levels for completeness). We set a couple of parameters in MaAsLin2 function such as: (**i**) cumulative sum scaling (CSS) normalization to raw count data to account for varying sequencing depth across samples, (**ii**) a negative binomial regression model for association testing (suitable for overdispersed count or relative abundance data), (**iii**) minimum abundance and prevalence thresholds to filter out very rare taxa (features present in at least 10% of samples and relative abundance >= 0.001 were required), and (**iv**) no additional transformation of the normalized data (setting transform = “NONE”).. Each model tested the association between a given taxon’s abundance (predictor of interest) and the global Aβ PET SUVR (outcome), including the covariates listed above. In formula form, for each genus:

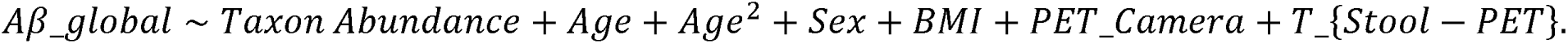

We also ran analogous models for each regional *A*β*_SUVR* in place of the *A*β*_global* term to see if certain brain regions drove the global associations. **Multiple-testing correction:** Given the number of taxa tested, we applied the Benjamini–Hochberg false discovery rate (FDR) correction to the *p*-values from these models. Associations with FDR-adjusted *p* (*q*-value) < 0.05 were deemed statistically significant.

### Sensitivity Analyses by *APOE* Genotype

We conducted a stratified analysis to examine whether gut-Aβ associations differ in *APOE* ε4 carriers vs. non-carriers. Essentially, we repeated the MaAsLin2 modeling described above in two subsets: (1) participants with **no *APOE*** ε**4 allele** (i.e., ε2/ε2, ε2/ε3, or ε3/ε3 genotypes), and (2) participants with **at least one *APOE*** ε**4 allele** (ε2/ε4, ε3/ε4, or ε4/ε4). This stratification was motivated by prior reports that *APOE* ε4 carriers may have a distinct microbiome profile and by our interest in whether the microbiome-amyloid link is present even in individuals without a genetic risk. We hypothesized that even *APOE* ε4 non-carriers would show an association between low SCFA-producer abundance and high Aβ (indicating a general gut-Aβ phenomenon), whereas *APOE* ε4 carriers might show the same pattern, perhaps even more strongly (if *APOE* ε4 exacerbates gut dysbiosis in concert with amyloid). In these subgroup analyses, we used the same modeling approach and covariates. Due to the smaller sample sizes in each stratum (N = 162 non-carriers, N = 57 carriers), power was reduced; therefore, we report nominal *p*-values and focus on the consistency of directionality and magnitude of effects relative to the full sample. Heatmaps (**Figures 2** and **3**) were generated to summarize the significant associations within each subgroup (p < 0.05, FDR correction applied within each subgroup analysis).

**Figure 1.**
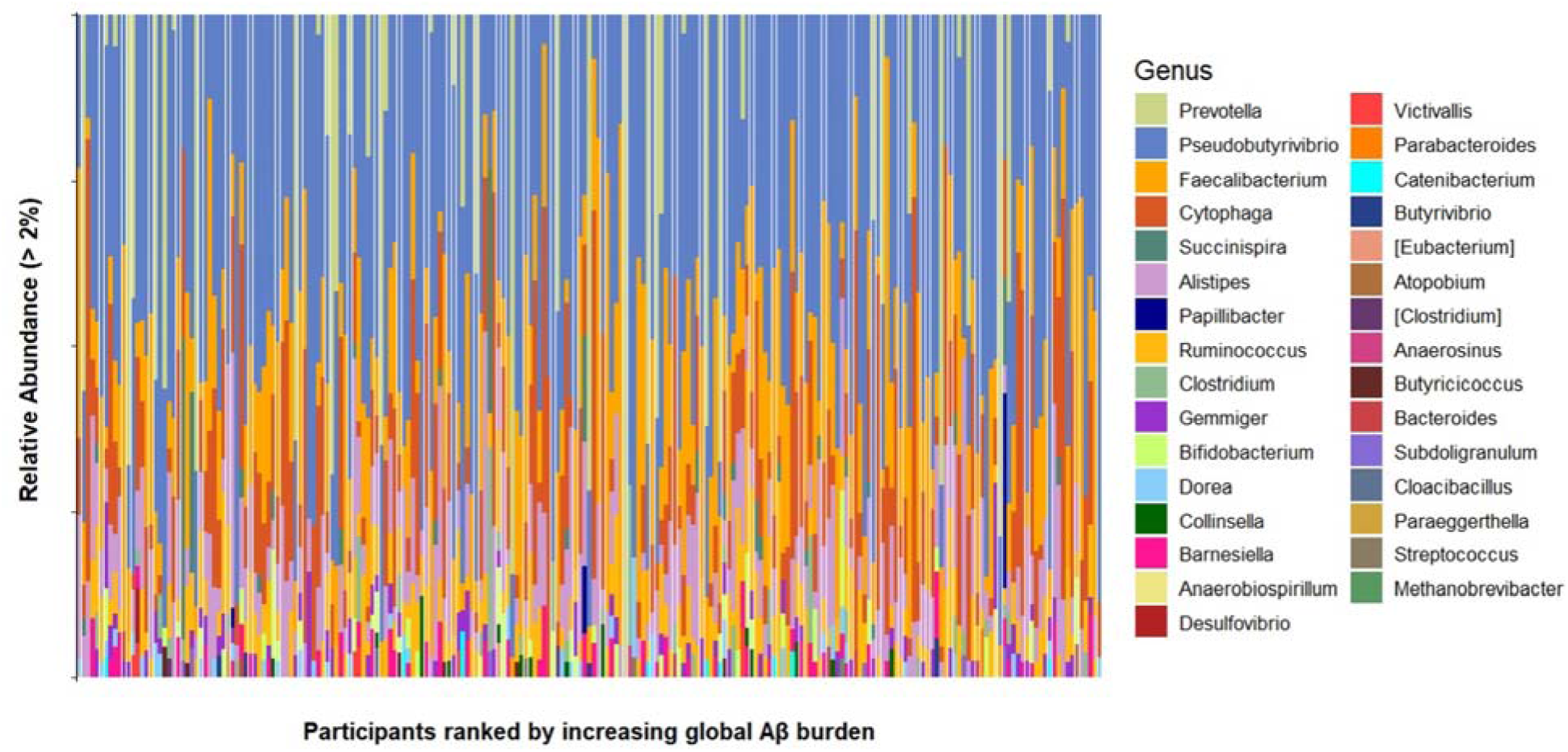
*Taxonomic profiling from stool samples of FHS subjects*. Stacked plot showing the microbiome profiles of predominant taxa in the stool samples of FHS individuals ordered from lowest to higher global Aβ burden at the genus level.

**Figure 2.**
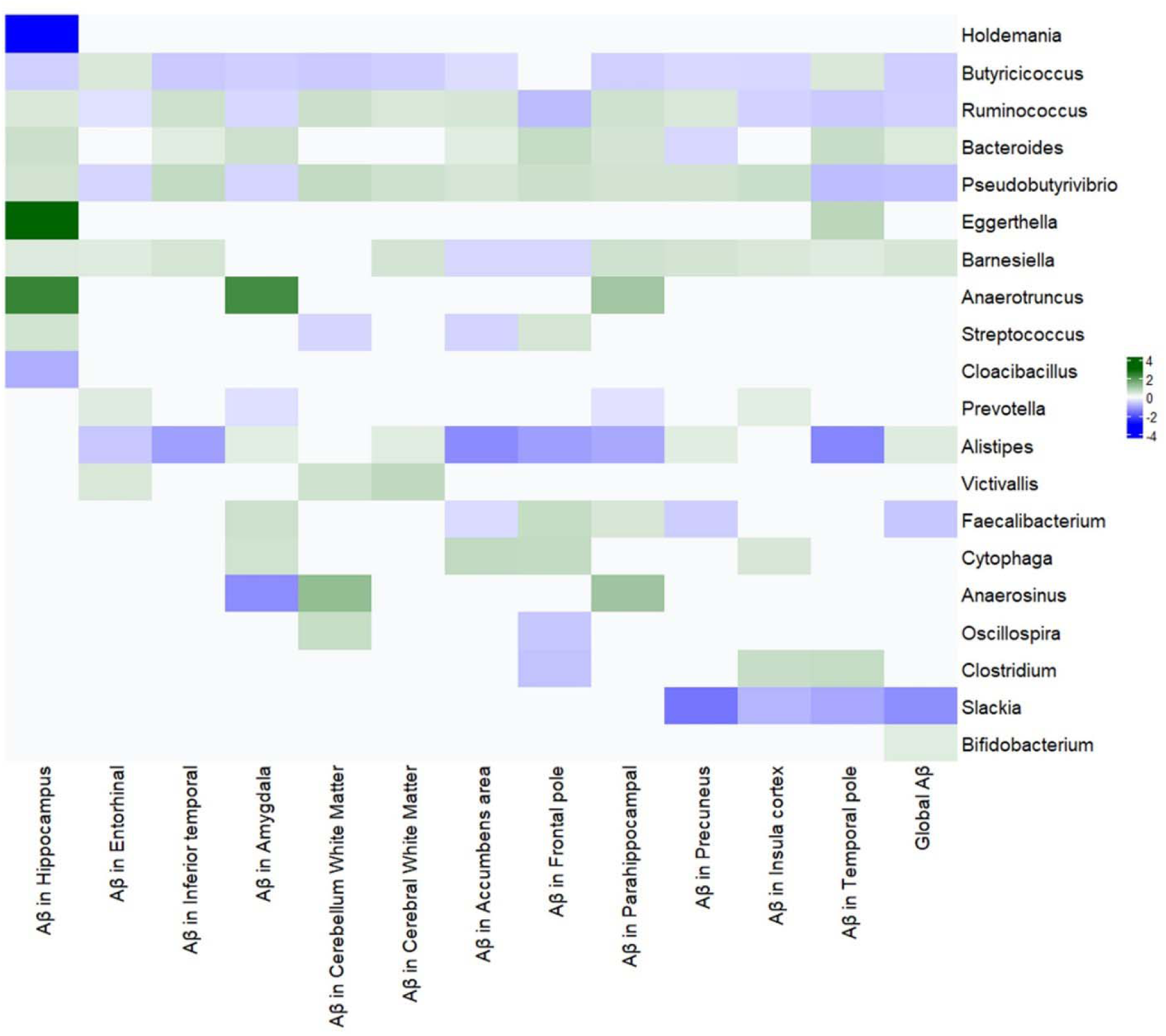
***Heatmap of multivariable associations between gut microbiome (genera) and amyloid-***β ***PET burden in the full sample***. Genera with BH-adjusted p<0.05 are shown. Warmer colors (green) indicate a positive association (higher abundance with higher Aβ), and cooler colors (blue) indicate a negative association (lower abundance with higher Aβ). The first column is global Aβ, followed by columns for regional Aβ measures (Accumbens, Amygdala, etc.). Notably, SCFA-producing genera (*Faecalibacterium*, *Ruminococcus*, *Butyricicoccus*, *Pseudobutyrivibrio*, *Slackia*) show negative associations (blue), while *Alistipes*, *Bacteroides*, *Barnesiella* show positive (green).

**Figure 3.**
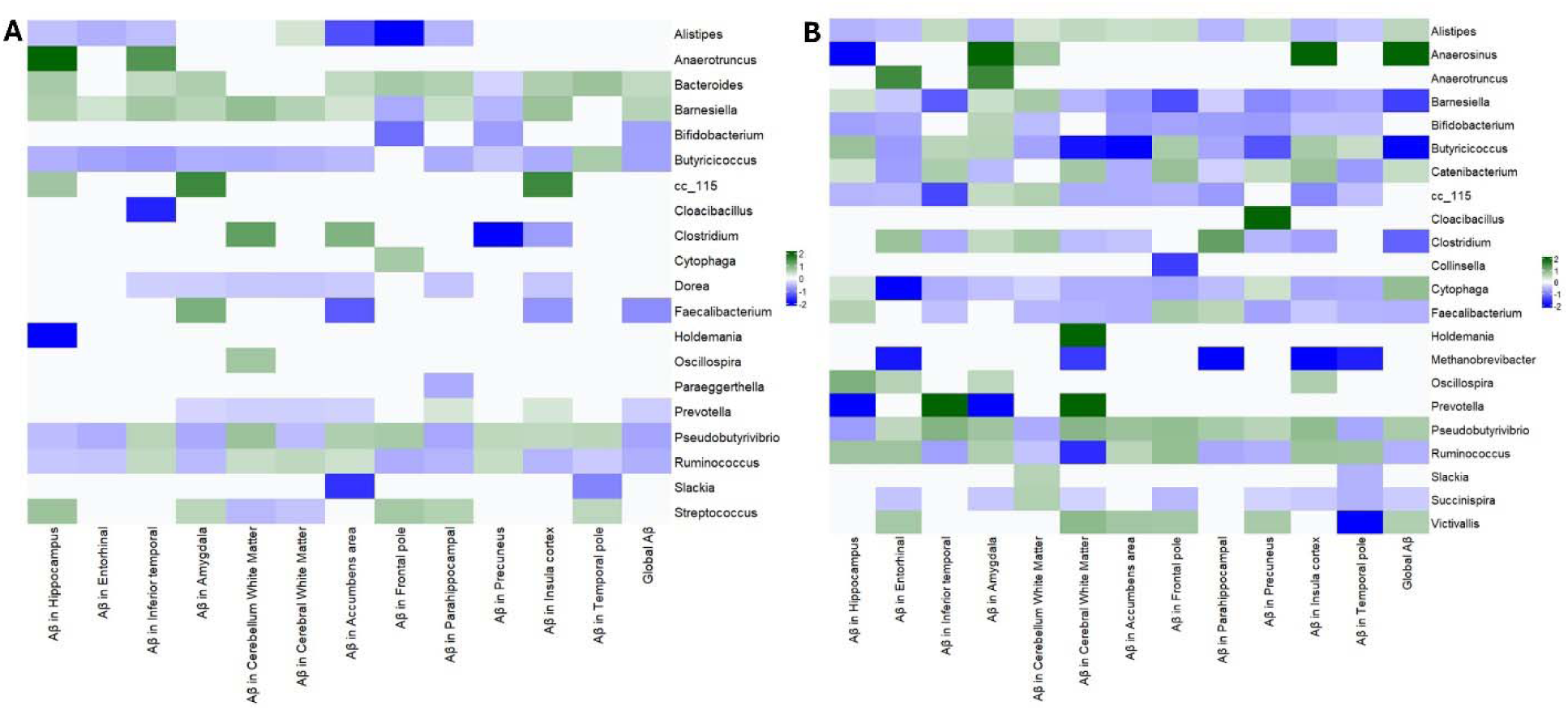
**(A) *Gut microbiome-A***β ***associations in APOE*** ε***4 non-carriers***. Heatmap similar to **Figure 2**, but for the subset of participants with no ε4 allele. The pattern is largely similar to the full sample (e.g., low *Faecalibacterium* with high Aβ). *Prevotella* appears as an additional negatively associated genus in this subgroup. **(B) *Gut microbiome–A***β ***associations in APOE*** ε***4 carriers***. Heatmap for participants with ≥1 ε4 allele. A broader array of associations is seen: in addition to those in Figure 2, *Barnesiella* and *Clostridium* are now negatively associated (blue) with Aβ, and several additional genera (bottom rows) are positively associated (green). This indicates a more pronounced dysbiosis in *APOE* ε4 carriers with high Aβ.

### Mediation Analysis

To formally test whether the gut microbiome mediates the relationship between *APOE* genotype and amyloid burden, we performed a causal mediation analysis using the mediation package in R.^63^ We focused on *APOE* ε4 carrier status (yes/no) as the exposure (X) and global Aβ PET

SUVR as the outcome (Y). We considered each candidate taxon that showed significant associations with Aβ in the earlier analyses as a potential mediator (M). For each such taxon, we set up two regression models:

- **Mediator model:** Taxon ∼ APOE ε4 + age + age^2^ + sex + BMI
- **Outcome model:** Aβ ∼ APOE ε4 + M + age + age^2^ + sex + BMI

All models were either linear or generalized linear, as appropriate (taxon abundances were modeled using negative binomial or an appropriate transformation). We then used the mediate function of the R package “*mediation*”^64^ to estimate the ***indirect effect*** of *APOE* ε4 on Aβ through the taxon (i.e., the mediation effect), as well as the direct effect not through the taxon. We employed nonparametric bootstrap resampling (1000 simulations) to compute confidence intervals for the mediated effect. An **indirect (mediated) effect** was considered statistically significant if the bootstrap *p*<0.05. We included multiple taxa in parallel only qualitatively (since formal multiple-mediator models are complex); instead, we tested each mediator separately. In reporting results, we focus on taxa that met the significance threshold for mediation.

All statistical tests were two-tailed, with an alpha level of 0.05 for significance (or FDR q<0.05 for multiple comparisons, as noted). Analyses were primarily descriptive/exploratory given the novelty of this area; thus, we also report trends (p<0.05) that align with hypotheses for completeness. Data analysis scripts and code for reproducing the results will be made available in a public repository upon publication.

## Results

### Participant Characteristics and Amyloid Burden

Relevant demographic and clinical characteristics of the studied cohort are summarized in **Table 1**. The mean age was 56.5 ± 8.3 years (range 32–73), and 58% were female. By design, all participants were free of clinical dementia, and overall had low comorbidity burdens (e.g., 8.8% with a history of diabetes, 3.1% with cardiovascular disease). *APOE* genotyping was successfully performed for 219 of the 227 participants (96.5%); the distribution of *APOE* genotypes in our sample was: 0.5% ε2/ε2, 10.9% ε2/ε3, 62.6% ε3/ε3, 3.2% ε2/ε4, 21.0% ε3/ε4, and 1.8% ε4/ε4. For analytical purposes, we classified individuals as ***APOE*** ε**4 carriers** (those with at least one ε4 allele; n = 57, 26%) or **non-carriers** (those with no ε4 allele; n = 162, 74%), consistent with common practice in stratifying *APOE*-related risk groups.

**Table 1.** Baseline characteristics of the study participants stratified by APOE ε4 status and summary of global/regional Aβ-PET values.

| Variables | Overall (N = 227) |
| --- | --- |
| Age, years | $56.5 \pm 8.3$ |
| Female, n (%) | 131 (58) |
| Camera, n (%) |  |
| Discovery GE-Smooth | 66 (29) |
| HR+ | 161 (71) |
| Time interval between stool collection and PET amyloid scans, weeks, median [Q1, Q3] | 110.0 [47.9, 239.6] |
| **Cardiovascular risk factors** |  |
| Triglycerides, mg/dL | $112.6 \pm 77.2$ |
| Systolic blood pressure, mmHg | $120 \pm 13$ |
| Diastolic blood pressure, mmHg | $76 \pm 8$ |
| Total cholesterol, mg/dL | $189.7 \pm 34.5$ |
| HDL cholesterol, mg/dL | $61.4 \pm 19.1$ |
| Treatment for hypertension, n (%) | 58 (25.55) |
| Stage I hypertension, n (%) | 70 (30.84) |
| Prevalent of cardiovascular disease, n (%) | 7 (3.08) |
| Current smoking, n (%) | 9 (3.96) |
| History of Diabetes, n (%) | 20 (8.81) |
| Body mass index, kg/m<sup>2</sup>, median [Q1, Q3] | 27.5 [24.4, 30.6] |
| **Global and Regional Aβ** |  |
| Global Aβ-PET | 1.08 ± 0.09 |
| Aβ-PET in the accumbens area | 1.09 ± 0.13 |
| Aβ-PET in the amygdala | 1.11 ± 0.08 |
| Aβ-PET in the cerebellum white matter | 1.42 ± 0.08 |
| Aβ-PET in the cerebral white matter | 1.36 ± 0.09 |
| Aβ-PET in the entorhinal cortex | 0.97 ± 0.07 |
| Aβ-PET in the frontal pole | 0.78 ± 0.18 |
| Aβ-PET in the hippocampus | 1.13 ± 0.08 |
| Aβ-PET in the inferior temporal cortex | 1.03 ± 0.11 |
| Aβ-PET in the insula cortex | 1.13 ± 0.08 |
| Aβ-PET in the parahippocampal cortex | 1.01 ± 0.08 |
| Aβ-PET in the precuneus cortex | 1.17 ± 0.13 |
| Aβ-PET in the temporal pole | 0.87 ± 0.07 |
| *APOE*, n (%) | N = 219 |
| $\epsilon 2/\epsilon 2$ | 1 (0.5) |
| $\epsilon 2/\epsilon 3$ | 24 (10.9) |
| $\epsilon 3/\epsilon 3$ | 137 (62.6) |
| $\epsilon 2/\epsilon 4$ | 7 (3.2) |
| $\epsilon 3/\epsilon 4$ | 46 (21.0) |
| $\epsilon 4/\epsilon 4$ | 4 (1.8) |

The global Aβ PET SUVR ranged from ∼0.9 to 1.3 (mean ± SD ≈ 1.08 ± 0.09).^65^ Approximately 23% of participants had high Aβ burden (above a typical positivity cut-off of ∼1.10 SUVR), consistent with the age range and inclusion of preclinical AD cases. Regional Aβ PET values demonstrated the expected pattern of amyloid accumulation: highest retention in frontal and cingulate regions (e.g., frontal pole SUVR 0.78 ± 0.18; precuneus 1.17 ± 0.13; cingulate included in composite) and relatively lower in medial temporal regions (entorhinal cortex 0.97 ± 0.07) and the hippocampus (1.13 ± 0.08) which often have delayed amyloid involvement. Notably, white matter regions (cerebral and cerebellar white matter) showed higher PiB uptake (as they are used for off-target binding references). These values align with established regional patterns of amyloid deposition.^66,67^

Out of 219 genotyped individuals, 26% carried at least one *APOE* ε4 allele. By definition, *APOE* ε4 carriers are at elevated risk for AD; accordingly, they had slightly higher mean global Aβ (1.11 ± 0.10) than non-carriers (1.07 ± 0.08), and a greater proportion of amyloid-positive cases (though many ε4 carriers were still amyloid-negative, given their middle age). *APOE* ε2 carriers (including ε2/ε2 and ε2/ε3) tended to have lower Aβ levels, though numbers were small.

Microbiome sequencing and processing were performed as described above. The relative abundance of the predominant genera in the stool samples, ranked from lowest to highest burden of global Aβ, is shown in **Figure 1**. The minimum read was 13,549. Across all samples, we found a median sequencing depth of 67,389 (interquartile, 50,348-77,626) paired-end reads. A total of ∼5,118 ASVs, agglomerated into 14 phyla, 70 families, and more than 140 genera, were shared across depths. The predominant genera were *Pseudobutyrivibrio*, *Faecalibacterium*, *Prevotella*, *Cytophaga*, *Ruminococcus*, and *Alistipes*. The processed data were used to assess the link between gut microbiome composition and Aβ burden in the brain.

### Gut Microbiome Composition and Global A**β** Burden (Full Sample Analysis)

We first examined the association between gut bacterial genera and global brain Aβ burden in the full sample (N = 227). Multivariable MaAsLin2 regression identified a set of microbial taxa significantly associated with global Aβ PET, after adjusting for covariates and correcting for multiple tests. **Figure 2** summarizes the significant associations at the genus level with global Aβ (and shows associations with regional Aβ for those genera). **Table 2** provides a full listing of all tested genera and their regression coefficients.

**Table 2.**
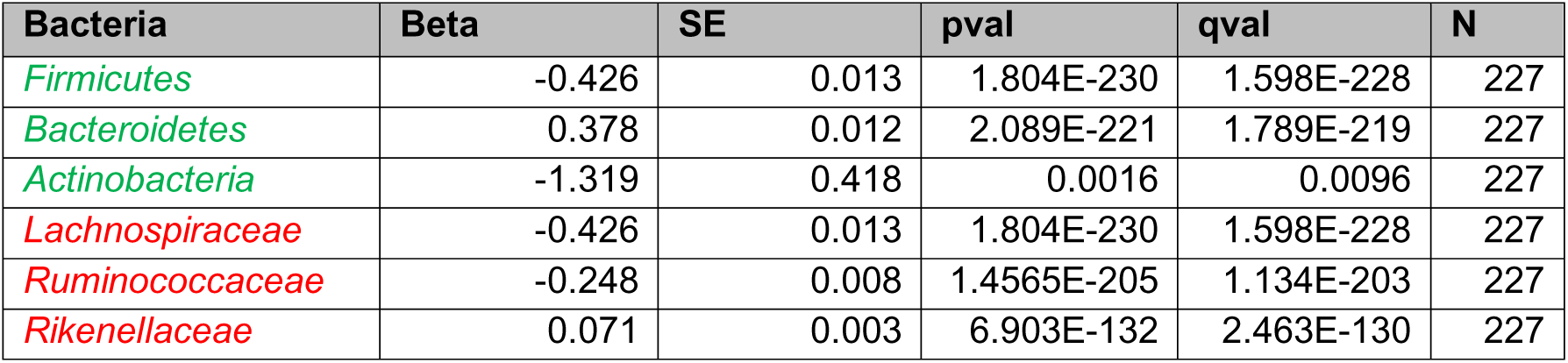
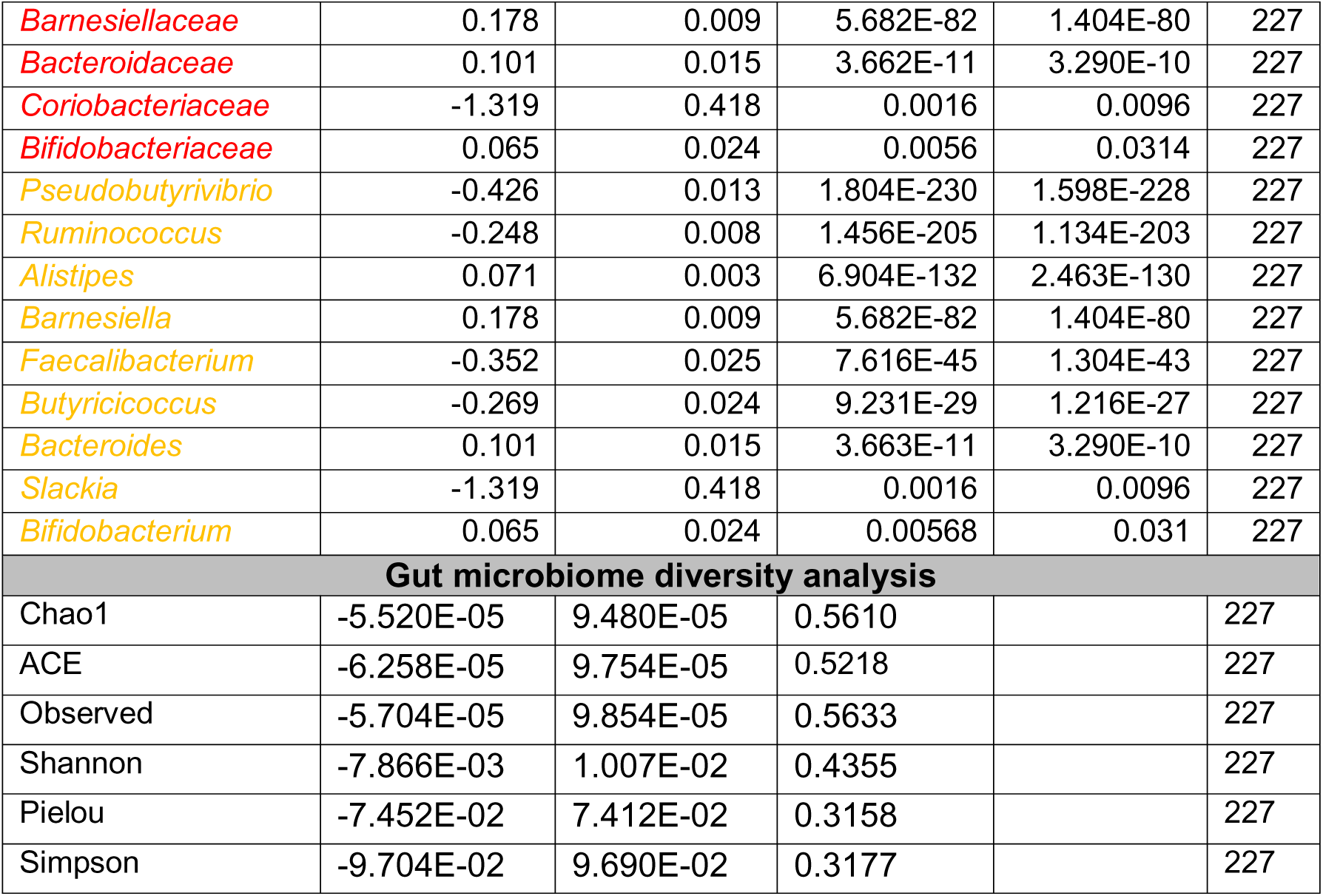
List of bacteria that showed association with global Aβ at the phylum (green), family (red), and genus (orange) levels. Multivariable association between global Aβ and gut microbial diversity using the model described in the Methods section.

### Key findings (global Aβ associations)

Higher global Aβ was associated with a lower relative abundance of several protective genera. In particular, genera in the family *Ruminococcaceae* were prominently implicated: *Faecalibacterium*, *Butyricicoccus*, and *Ruminococcus* all showed significant negative associations with global Aβ (β (SE) ≈-0.35 (0.025),-0.27 (0.024),-0.25 (0.008), respectively, all *q*<0.001). Indicating that a standard deviation unit (1-SDU) greater abundance of *Faecalibacterium*, *Butyricicoccus*, and *Ruminococcus* was associated with a 0.35, 0.27, and 0.25 lower global Aβ burden (by SDU), respectively (**Figure 2**, **Table 2**).. These bacteria are known to harbor anti-inflammatory properties in the gut and can produce SCFAs. Similarly, *Pseudobutyrivibrio* belonging to family *Lachnospiraceae* (another protective bacterium) was significantly reduced in those with higher Aβ levels (β (SE) ≈-0.42 (0.013), suggesting a 0.42 lower global Aβ burden for an SDU increase in the abundance of *Pseudobutyrivibrio*). Another genus, *Slackia* (a member of the family *Coriobacteriaceae* that produces acetate), was also less abundant, with a greater Aβ (**Figure 2**, blue cells indicating a negative correlation). The associations were strong, indicating an extremely robust relationship. These results suggest a pattern of SCFA depletion in the gut microbiomes of individuals with elevated brain amyloid. In these models, β coefficients indicate the difference in the Aβ burden measures (by SDU) associated with a 1-SDU higher abundance of a specific bacterium.

Conversely, higher global Aβ was associated with an increased abundance of certain genera often considered opportunistic or pro-inflammatory. Notably, *Alistipes* (order *Bacteroidales*) and *Bacteroides* (family *Bacteroidaceae*) were positively associated with Aβ burden (β ≈ 0.07, 0.10, *q*<0.01). Both genera are Gram-negative and can produce endotoxins (lipopolysaccharides); *Alistipes,* in particular, has been linked to pro-inflammatory states. Additionally, *Barnesiella* (family *Barnesiellaceae*) was more abundant in those with higher Aβ (*q*<0.001). Interestingly, *Barnesiella* is typically considered to have beneficial effects on gut health (e.g., it can protect against *C. difficile* infection), but in our data, it positively correlated with the amyloid burden. Finally, *Bifidobacterium* (a genus often considered beneficial and probiotic) showed a modest positive association with Aβ (β (SE) ≈ +0.07 (0.024), *q*=0.03). The *Bifidobacterium* finding was somewhat unexpected, as we anticipated it to be lower in high-Aβ individuals; we revisit this in subgroup analyses. In summary, the global Aβ burden was associated with a gut bacterial profile indicative of dysbiosis, characterized by a reduction in SCFA-producing, anti-inflammatory microbes and an enrichment of bacteria that could contribute to inflammation or thrive in inflammatory environments.

At higher taxonomic levels, we observed analogous patterns (**Table 2**). For instance, the family *Ruminococcaceae* (which includes *Faecalibacterium* and *Ruminococcus*) was found to be lower in participants with high-Aβ globally, and the family *Lachnospiraceae* (which includes *Butyricicoccus* and *Pseudobutyrivibrio*) was also decreased, although the *Lachnospiraceae* family overall did not reach significance after correction. *Coriobacteriaceae* (family of *Slackia*) was negatively associated with Aβ. On the other hand, the families *Bacteroidaceae*, *Barnesiellaceae*, and *Rikenellaceae* (the latter includes *Alistipes*) were positively associated with Aβ (all *q*<0.05). At the broad phylum level, we noted a trend that high-Aβ subjects had a higher *Bacteroidetes*:*Firmicutes* ratio compared to low-Aβ subjects. Phylum *Bacteroidetes* was relatively increased with greater Aβ, whereas phylum *Firmicutes* (which encompasses most SCFA producers) was reduced (**Table 2**). There was also an apparent lower proportion of *Actinobacteria* (which includes *Bifidobacteria*) in high-Aβ individuals, although the *Bifidobacterium* genus itself was slightly higher, as noted (suggesting other *Actinobacteria* genera were lower). These broad patterns align with a pro-inflammatory microbiome state in individuals with higher amyloid levels.

In addition, to evaluate whether gut microbiota diversity is associated with Aβ loads, we conducted a multivariable linear regression analysis after controlling for the covariates indicated in the methods section. We found that the Shannon index (β ≈-0.008, *p* = 0.43) and Pielou evenness (β ≈-0.074, *p* = 0.31) showed negative relationships that were not statistically significant with Aβ burden (**Table 2**). Supporting the evidence that there is no significant difference in gut microbial diversity in individuals with higher Aβ vs. lower Aβ levels on average.

### Regional A**β** associations mirror global trends

To understand whether the global Aβ-microbiome associations were driven by particular brain regions, we examined associations with regional Aβ PET measures. We found that, in general, the same genera associated with global Aβ were also associated with Aβ in multiple individual regions (**Figure 2** shows a heatmap across regions for each genus). For example, *Ruminococcus* was negatively associated not only with global Aβ but also with Aβ in the entorhinal cortex, amygdala, frontal pole, insular cortex, and temporal pole (all *q*<0.01). This suggests that amyloid in those specific regions may be more susceptible to changes in gut microbial composition. *Faecalibacterium*’s negative association with global Aβ was also observed with Aβ in the nucleus accumbens area and precuneus, two regions known to show early amyloid deposition. *Butyricicoccus* was inversely associated with Aβ in most regions, except for a few (the entorhinal, temporal pole, and frontal pole did not reach significance for *Butyricicoccus*, although the directions were consistent). *Pseudobutyrivibrio* showed a remarkably consistent negative association with Aβ across many regions (particularly entorhinal, amygdala, and temporal pole). *Slackia* was negatively associated with Aβ in the precuneus, insula, and temporal pole as well. On the “increased” side, *Bacteroides*’ positive link with global Aβ was reflected in its association with Aβ in a broad set of regions: hippocampus, inferior temporal, amygdala, accumbens, frontal pole, parahippocampal, and temporal pole (all *q*<0.05). *Barnesiella*’s positive association was consistent across most regions, except for a few, where it was not significant (in the accumbens, frontal pole, amygdala, or cerebellar white matter). In sum, regional amyloid measures generally reinforced the global findings, indicating that the gut microbiome correlations are not driven by a single anomalous brain region but rather reflect a whole-brain amyloid load effect. Regions known to accumulate amyloid early (like precuneus, frontal cortex, and temporal cortex) showed particularly strong microbiome ties. To assess the degree of correlation between Aβ loads in different brain regions and global Aβ, we performed a Spearman correlation analysis. All the regional Aβ burdens and global Aβ exhibited significant positive correlation with correlation coefficients ranging between 0.24 and 0.75.

These results bolster our confidence that the microbiome-Aβ relationships are biologically meaningful. The consistency across brain regions and the coherence of the bacteria involved (mostly SCFA producers and a few prominent *Bacteroidetes* genera) suggest a scenario in which a higher brain amyloid burden is associated with deficient gut microbial production of beneficial metabolites and a relative overrepresentation of potentially pro-inflammatory microbes. This set the stage for exploring whether the *APOE* genotype influences these associations.

### *APOE* Genotype Modifies the Gut–A**β** Association Patterns

We next performed stratified analyses to determine how the presence or absence of the *APOE* ε*4* allele might alter the relationship between gut microbiome and amyloid burden. We separated participants into *APOE* ε4 **non-carriers** (N = 162, with no ε4 allele) and *APOE* ε4 **carriers** (N = 57, with at least one ε4 allele). While the limited sample size in the carrier group cautions against over-interpretation, several interesting patterns emerged.

### Gut microbiome and A**β** in *APOE* **ε**4 non-carriers

Even among individuals without *APOE* ε4, we observed that higher Aβ levels were associated with a microbiome profile characterized by reduced SCFA-producing bacteria, a pattern similar to that observed in the full sample. Specifically, *Faecalibacterium*, *Butyricicoccus*, *Ruminococcus*, and *Pseudobutyrivibrio* were all significantly negatively associated with global Aβ in APOE ε4 non-carriers (all *p*<0.01; **Figure 3A**). This suggests that the link between Aβ deposition and depletion of these beneficial gut bacteria is a general phenomenon, not solely driven by *APOE* ε4. Additionally, in *APOE* ε4 non-carriers, *Barnesiella* and *Bacteroides* remained positively associated with global Aβ (mirroring the full sample).

However, there were a few notable differences in the non-carrier group relative to the full cohort. In *APOE* ε4 non-carriers, *Bifidobacterium* showed a negative association with Aβ (opposite to the slight positive trend seen in the full sample). That is, non-carriers with high Aβ had lower *Bifidobacterium* abundance, which aligns with expectations since *Bifidobacterium* is generally beneficial. This discrepancy suggests that the small positive association we saw earlier for *Bifidobacterium* might have been driven by the *APOE* ε4 carrier subset. Indeed, we will see that in *APOE* ε4 carriers, *Bifidobacterium* was not significantly associated with Aβ (and possibly trended positive, diluting the overall effect). Also, in *APOE* ε4 non-carriers, *Prevotella* (genus in family *Prevotellaceae*) emerged as significantly negatively associated with global Aβ (*q*<0.05), whereas in the full sample *Prevotella* had shown no significant relationship. *Prevotella* is a fiber-degrading bacterium that can produce propionate; a lower *Prevotella* in high-Aβ non-carriers could indicate another SCFA-related deficit. Meanwhile, *Alistipes* and *Slackia* were not significantly associated with Aβ in non-carriers (though directions were consistent with the full sample). At the family level, the patterns for *Ruminococcaceae*, *Lachnospiraceae*, *Coriobacteriaceae*, *Barnesiellaceae*, and *Bacteroidaceae* (**Figure 4A**) in non-carriers were largely the same as in the full cohort.

**Figure 4.**
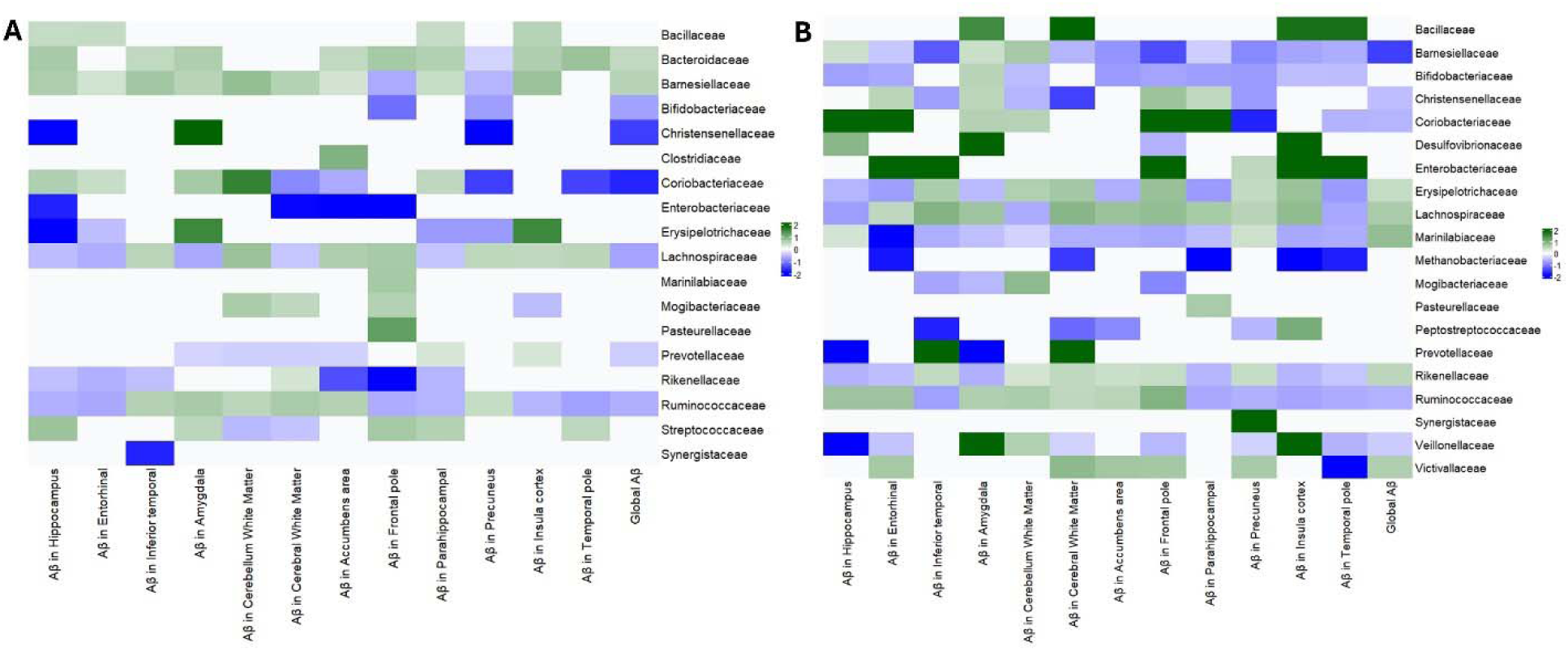
**(A) *Gut microbiome-A***β ***associations in APOE*** ε***4 non-carriers***. Findings close to those reported in the full sample. **(B) *Gut microbiome–A***β ***associations in APOE*** ε***4 carriers***. Heatmap for participants with ≥1 ε4 allele. A broader array of associations is seen.

In summary, *APOE* ε4 non-carriers exhibited the same core gut-Aβ associations: lower levels of anti-inflammatory bacteria, higher *levels of Bacteroides/Barnesiella, and* higher Aβ. They also showed that *Bifidobacterium* and *Prevotella* (both generally health-associated genera) were inversely linked to Aβ in this subgroup, a signal that might have been masked in the combined analysis. This consistency in non-carriers underscores that the gut microbiome’s relationship with amyloid is not contingent on *APOE* ε4 status; even without the high-risk allele, having a more “dysbiotic” microbiome correlates with greater amyloid deposition.

### Gut microbiome and A**β** in APOE **ε**4 carriers

We then assessed the associations in *APOE* ε4 carriers, who constitute the higher-risk group for AD. Notably, *APOE* ε4 carriers exhibited a more extensive array of significant microbiome associations with Aβ, both in the direction observed in non-carriers and with some additional taxa not previously identified (**Figure 3B**). This suggests that carrying an *APOE* ε4 allele may exacerbate the gut microbial changes associated with amyloid accumulation.

In *APOE* ε4 carriers, greater global Aβ was strongly associated with a lower abundance of SCFA-producing genera, including *Faecalibacterium* and *Ruminococcus* (again), but also lower *Barnesiella* and lower *Clostridium*. The finding that *Barnesiella* was *decreased* with high Aβ in *APOE* ε4 carriers is especially noteworthy because, in non-carriers (and overall), *Barnesiella* tended to increase with Aβ. This indicates a potential genotype interaction: *APOE* ε4 carriers with high Aβ levels lose *Barnesiella* (a generally beneficial commensal), whereas non-carriers with high Aβ levels do not show this loss. *Clostridium* (sensu stricto, a core anaerobic genus with many butyrate-producing species) was also significantly lower in high-Aβ *APOE* ε4 carriers but not significant in non-carriers. Additionally, *Succinispira* (a lesser-known genus that ferments succinate) was identified as reduced in high-Aβ *APOE* ε4 carriers.

On the other hand, APOE ε4 carriers with higher Aβ showed increases in certain genera that were not as evident in non-carriers. *Alistipes* and *Pseudobutyrivibrio* remained positively associated with Aβ (as before), and new associations emerged: *Cytophaga*, *Anaerosinus*, *Catenibacterium*, and *Victivallis* were all significantly more abundant with greater Aβ in *APOE* ε4 carriers (these genera did not reach significance in non-carriers). Many of these are relatively uncommon gut genera; for instance, *Catenibacterium* and *Anaerosinus* are anaerobes whose roles are not well-characterized, and *Victivallis* is a member of the phylum *Lentisphaerae*. Their increase might indicate a wider dysbiosis under *APOE* ε4 conditions, or perhaps these taxa fill niches when protective taxa dwindle.

Moreover, at the family level, *APOE* ε4 carriers exhibited broader alterations: families *Ruminococcaceae*, *Barnesiellaceae*, *Christensenellaceae*, *Peptostreptococcaceae*, and *Veillonellaceae* were all significantly lower in abundance, with higher Aβ levels (all *p* < 0.05). Notably, *Christensenellaceae* is a family often associated with a healthy gut and metabolic benefits; its reduction in high-Aβ *APOE* ε4 carriers is consistent with our narrative of protective taxa depletion. On the other hand, families such as *Rikenellaceae*, *Marinilabiliaceae*, *Erysipelotrichaceae*, *Lachnospiraceae*, *Bacillaceae*, and *Victivallaceae* exhibited higher abundance and greater Aβ levels in *APOE* ε4 carriers (**Figure 4B**). Several of these contain pro-inflammatory or less beneficial members. For example, *Erysipelotrichaceae* increases have been linked to metabolic syndrome and inflammation.

In essence, *APOE* ε4 carriers displayed a more pronounced “*amyloid-associated dysbiosis*” than non-carriers. The same beneficial bacteria (*Faecalibacterium*, *Ruminococcus*, etc.) were depleted, but additionally, *APOE* ε4 carriers lost other beneficial taxa (*Barnesiella*, *Christensenellaceae*, etc.) and experienced a bloom of various minor genera. This suggests an interaction where *APOE* ε4 status might exacerbate gut microbial responses to (or drivers of) amyloid accumulation. It is intriguing that *Barnesiella* transitions from a positive association in non-carriers to a negative one in carriers, possibly indicating that *APOE* ε4 carriers cannot maintain *Barnesiella* in the face of mounting amyloid or inflammation, whereas non-carriers can (hence, non-carriers paradoxically showed an increase in *Barnesiella* with Aβ, perhaps as a compensatory response). The lower *Christensenellaceae* in *APOE* ε4 carriers with high Aβ is also of interest, given *Christensenellaceae’s* links to low inflammation and healthy metabolism have been previously reported.

Taken together, our stratified analyses indicate that the gut microbiome-amyloid relationship exists irrespective of *APOE* genotype but is broadened and intensified in *APOE* ε4 carriers. Even *APOE* ε2/ε3 individuals exhibit the core pattern of anti-inflammatory taxa loss with amyloid, whereas *APOE* ε4 carriers show this pattern, along with additional dysbiotic shifts. This raises the possibility that the *APOE* ε4 genotype may be associated with gut dysbiosis, which could promote AD pathology. To further investigate causality, we next examined whether the gut microbiome might mediate the effect of *APOE* ε4 on amyloid burden.

### Mediating Role of the Gut Microbiome in APOE **ε**4–Related A**β** Burden

Finally, we conducted mediation analyses to determine whether certain gut bacteria significantly mediate the association between *APOE* ε4 carrier status and Aβ burden. In our data, carrying an APOE ε4 allele was associated with higher amyloid burden (linear model β = 0.055, SE = 0.013, *p* = 5.6×10^-5^ after adjusting for age, sex, and BMI). This confirms the known deleterious effect of *APOE* ε4 on amyloid accumulation in our cohort. We then examined whether this effect might be transmitted via changes in the gut microbiome.

Out of the set of candidates mediating taxa (those showing significant Aβ associations in both *APOE* strata), we identified two gut microbial taxa that exhibited a significant indirect mediation effect: the genera *Ruminococcus*, *Butyricicoccus*, and *Clostridium*, as well as the family *Christensenellaceae* (**Figure 5**). These bacteria mediated a portion of 0.07%, 0.13%, 0.06%, and 0.11% of the *APOE* ε4’s effect on Aβ (each with *p* < 0.05), respectively. In both cases, the mediation followed a plausible biological direction: *APOE* ε4 carriers tended to have a lower abundance of *Ruminococcus*, *Butyricicoccus*, *Clostridium*, and *Christensenellaceae* (path a: genotype → microbiome), and lower levels of these microbes were in turn associated with higher Aβ (path b: microbiome → amyloid), yielding a significant indirect effect of *APOE* ε4 via these microbes. In other words, part of the reason *APOE* ε4 carriers have more amyloid may be that *APOE* ε4 leads to a deficiency of these taxa, specifically SCFA-producing, inflammation-modulating bacteria. The reduction of these beneficial microbes could contribute to a pro-amyloid environment. The mediation analysis quantifies this: we estimated that a significant fraction of *APOE* ε4’s effect (on the order of 0.3–0.4%) on Aβ burden was mediated by *Ruminococcus*, *Butyricicoccus*, *Clostridium,* and *Christensenellaceae*, though these estimates have wide confidence intervals. While the mediating pathways exist (since indirect effect is statistically significant), the effects (strength of the indirect effect) appeared quite modest. One possible explanation is that each bacterium might account for a small portion of the link between *APOE* ε4 and amyloid-beta burden.

**Figure 5.**
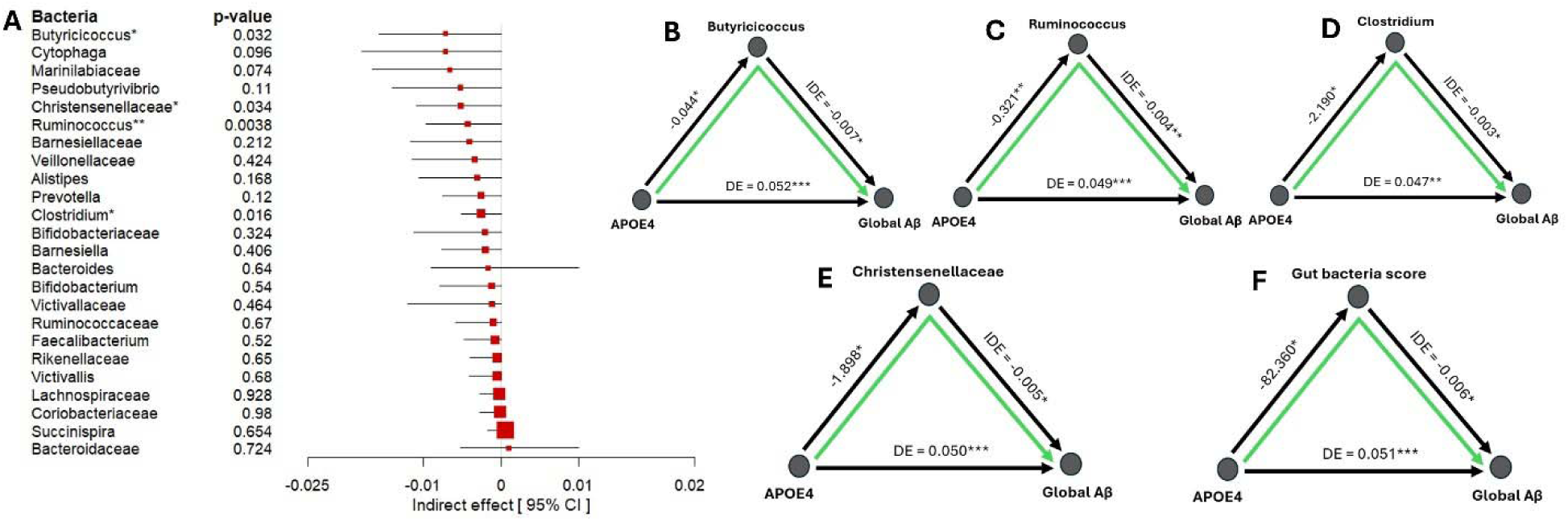
***Mediation analysis of gut microbiome in the APOE-A***β ***relationship***. **(A)** Indirect effect estimates for candidate mediating taxa. *Ruminococcus* and *Christensenellaceae* show significant mediation (indirect effect *p*<0.01), meaning they carry part of the APOE ε4 → Aβ effect. **(B, C, D, E)** Example mediation plots for *Ruminococcus*, *Butyricicoccus*, *Clostridium*, and *Christensenellaceae*, illustrating that APOE4 status is associated with lower abundance of these taxa, which in turn is associated with higher Aβ. (* – *p*<0.05, ** – *p*<0.01, *** – *p*<0.001 for mediation effect). **(F)** Mediation plot for gut bacteria score, demonstrating the linear combination effect of *Ruminococcus*, *Christensenellaceae*, *Butyricicoccus*, and *Clostridium* in the association between *APOE* ε4 and Aβ burden.

The majority of tested bacteria showed non-significant mediation effects, and notably, most of those effects were negative in sign. A negative indirect effect here would mean that *APOE* ε4 decreases the bacterium, which in turn increases Aβ (in the same direction as we observed for protective bacteria). So, while not individually significant, the consistent negative direction across many SCFA producers suggests a general mediating trend: *APOE* ε4 likely drives a broad reduction in beneficial gut microbes, collectively contributing to higher amyloid levels, even if only a couple reach statistical significance individually. Hypothesizing that bacteria may act in concert and have a combined effect on Aβ deposition, we sought to establish a model that encompasses all key bacteria related to Aβ in our cohort. We built a gut bacteria score based on a linear combination of bacteria demonstrating *p*<0.05 for the indirect effect. These bacteria include *Ruminococcus*, *Butyricicoccus*, *Clostridium*, and *Christensenellaceae*. Then, the model was constructed as follows: 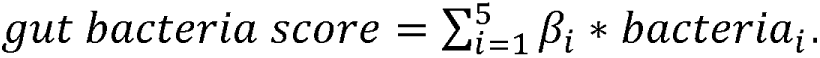 The results also revealed the mediating role of gut microbiome, supporting the hypothesis that a colony of bacteria may act together in the association between *APOE* genotype and Aβ burden (indirect effect =-0.006, 0.14% of *APOE*’s effect mediated by this gut bacteria score, *p*<0.05, **Figure 5F**).

### Predicted Functional Profile of Gut Microbiome in Relation to A**β**

Given that our microbiome data are based on 16S profiles, we utilized predictive metagenomic tools (PICRUSt2 via the ggpicrust R framework^68^) to infer the functional capacity of the gut microbiome, specifically regarding SCFA metabolism. We identified KEGG orthologs **(KOs)** in microbial genomes that are involved in the synthesis of SCFAs and tested which of these predicted functions were associated with global Aβ levels.

From ∼10,473 KOs predicted by PICRUSt, we curated a list of 299 KOs that are functionally annotated as related to SCFA synthesis (covering pathways for acetate, propionate, butyrate, succinate, lactate, pyruvate fermentation, etc.). We then ran MaAsLin2 regression for each KO (similar to the taxon models) to determine if KO abundance was associated with global Aβ, while adjusting for the same covariates. Results were visualized in a volcano plot (**Figure 6** and Table 3) and summarized by pathway (**Figure 7**).

**Figure 6.**
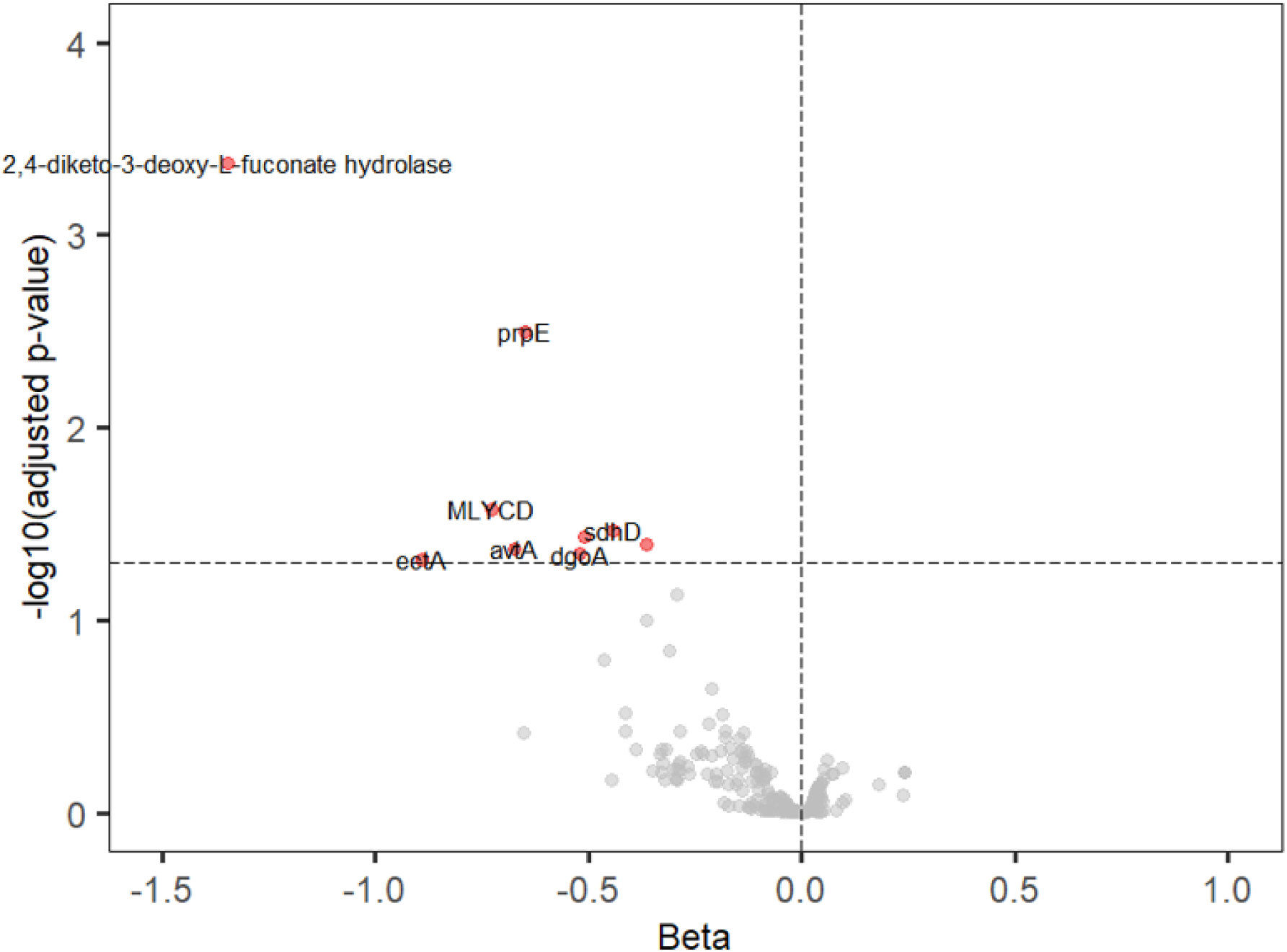
***Volcano plot of associations between predicted microbial gene functions (KEGG orthologs) and global A***β. Each point is a KO (SCFA-related KOs in color). The x-axis is the regression coefficient (effect of high Aβ on KO abundance), and the y-axis is –log10(*p*-value). KOs to the left (negative direction) are lower in high-A individuals. Nine SCFA-related KOs (red diamonds) surpass the significance threshold (dashed line corresponds t FDR q=0.05), all with negative coefficients, meaning they are significantly depleted with greater Aβ. No SCFA-related KO appears on the right side as significantly increased. This suggests a targeted loss of SCFA production genes in the high-amyloid microbiome.

**Figure 7.**
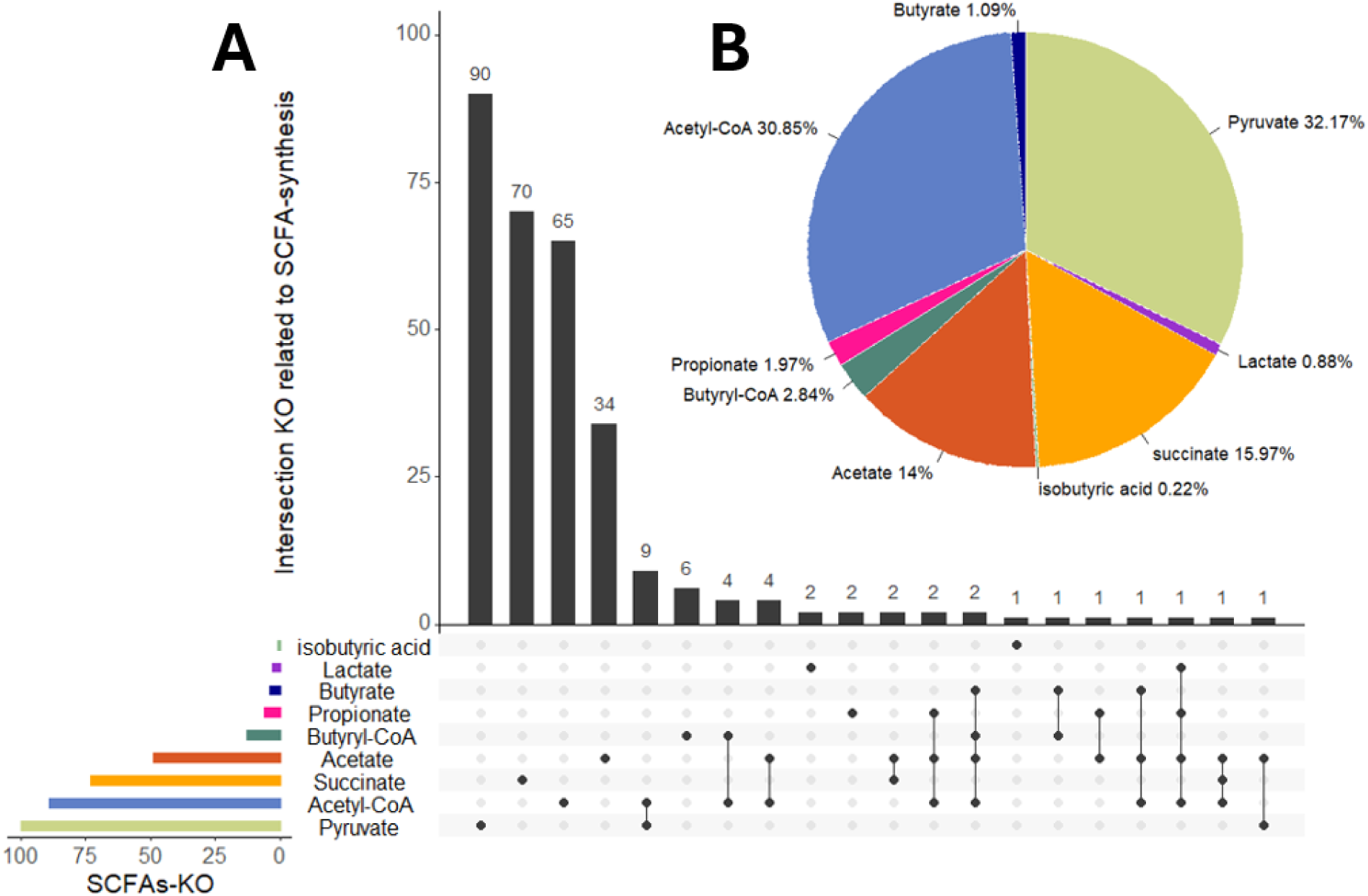
***Predicted functional potential of gut microbiome related to SCFA metabolism***. Bar charts showing th distribution of 299 SCFA-related KEGG orthologs (KOs) across major metabolic pathways (acetate, butyrate, propionate, succinate, etc.). Highlighted in red are pathways where multiple KOs were significantly less abundant in high-Aβ individuals. For example, several enzymes in the pyruvate→acetyl-CoA→butyrate pathway and in the propionate fermentation pathway were under-represented with higher Aβ, indicating a systemic reduction in SCFA biosynthetic capacity.

**Table 3.** List of KEGG orthologues that showed significant association with global Aβ burden.

| ID | Description | beta | stderr | pval | qval | N |
| --- | --- | --- | --- | --- | --- | --- |
| K18336 | 2,4-diketo-3-deoxy-L-fuconate hydrolase | -1.347 | 0.284 | 2.038E-06 | 0.000419 | 227 |
| K01908 | propionyl-CoA synthetase (prpE) | -0.649 | 0.153 | 2.199E-05 | 0.003167 | 227 |
| K01578 | malonyl-CoA decarboxylase (MLYCD) | -0.726 | 0.206 | 0.00042 | 0.026402 | 227 |
| K00242 | succinate dehydrogenase subunit D (sdhD) | -0.442 | 0.129 | 0.00059 | 0.034114 | 227 |
| K10764 | homoserine O-acetyltransferase (metZ) | -0.509 | 0.152 | 0.00082 | 0.036576 | 227 |
| K00163 | alanine transaminase (avtA) | -0.363 | 0.116 | 0.00169 | 0.040186 | 227 |
| K00835 | pyruvate dehydrogenase E1 component (aceE) | -0.673 | 0.119 | 0.00168 | 0.042186 | 227 |
| K01631 | 3-dehydrogalactonate aldolase (dgoA) | -0.521 | 0.118 | 0.00173 | 0.045186 | 227 |
| K06718 | ectoine synthase (ectA) | -0.892 | 0.130 | 0.00262 | 0.048180 | 227 |

We found nine SCFA-related microbial genes (KOs) that were significantly less abundant in individuals with higher global Aβ burden (BH-adjusted *p* < 0.05 for each; no KO was higher with high Aβ after FDR correction). These nine KOs correspond to enzymes in various metabolic pathways, largely consistent with reduced SCFA production potential in high-Aβ microbiomes.

All these KOs were significantly under-represented in the gut microbiomes of participants with higher Aβ (with effect sizes indicating ∼20–30% lower abundance per unit increase in Aβ SUVR, on average). No SCFA-related KO was found at higher levels in high-Aβ individuals, suggesting a global functional deficit in SCFA production capacity. **Figure 6** (volcano plot) highlights these nine KOs among the many tested (with most KOs showing no association, as expected under null). **Figure 7** provides a more intuitive overview: it shows the distribution of all SCFA-related KOs across major SCFA pathways (pyruvate to acetate, succinate to propionate, etc.), illustrating that the pathways for pyruvate/acetyl-CoA conversion, propionate biosynthesis, and succinate utilization had multiple KOs significantly depleted with high Aβ.

In practical terms, these functional predictions align with our taxonomic findings. For instance, the reduction in *Ruminococcaceae* and *Lachnospiraceae* (butyrate producers) in high-Aβ subjects is reflected by lower gene abundance in pyruvate to butyrate conversion (e.g., pyruvate dehydrogenase, which feeds acetyl-CoA for butyrate synthesis, was lower). Similarly, lower *Prevotellaceae* and other propionate producers correspond with lower propionyl-CoA synthetase. Although these predictions need validation with shotgun metagenomics or metabolite profiling, they provide supporting evidence that microbial metabolic functions related to SCFA production are suppressed in individuals with greater amyloid burden.

To sum up, the functional inference suggests that the gut microbiomes of high-amyloid individuals are not only taxonomically shifted but also functionally impaired in producing neuroprotective metabolites, such as butyrate and propionate. This could result in lower circulating SCFA levels reaching the brain, potentially weakening beneficial effects such as maintenance of the blood-brain barrier and suppression of neuroinflammation. It strengthens the case that SCFA depletion is a mechanism by which gut dysbiosis might influence AD pathology.

## Discussion

Our study reveals a new connection between host genetic variation (specifically, *APOE* genotype) and the gut microbiome, which is associated with Aβ deposits—a key feature of Alzheimer’s disease. Using a well-phenotyped cohort from the Framingham Heart Study, we found that individuals with higher brain amyloid-β levels have a distinct gut microbiome, characterized by a deficiency in bacteria that provide protective benefits and are known to produce short-chain fatty acids (SCFAs). This microbiome profile is also associated with the *APOE* genotype. To our knowledge, only a handful of human studies have shown a direct link between *APOE* status, the gut microbiome, and brain amyloid deposition. These findings shed light on the gut-brain axis as a potential mediator of genetic risk for AD and suggest that modulating the gut microbiota could be a novel avenue for mitigating *APOE*-related AD risk.

### Summary of key findings

We found that higher global Aβ load was associated with a decrease in SCFA-producing gut bacteria, most notably members of the *Ruminococcaceae* family (*Ruminococcus*, *Faecalibacterium*) and some *Lachnospiraceae* (e.g., *Butyricicoccus*, *Pseudobutyrivibrio*). Many of these genera play a major role in protecting the colon. This microbiome signature, which can be viewed as a loss of “good” commensals, is important because SCFA-producing bacteria are key microbes with anti-inflammatory and neuroprotective effects. The loss of protective taxa we observe in high-amyloid individuals is consistent with a pro-inflammatory gut environment. Indeed, depletion of *Ruminococcaceae* has been causally linked to inflammatory responses in prior studies.^69,70^ For example, *Faecalibacterium prausnitzii* (a *Ruminococcaceae* member) is known to produce butyrate and anti-inflammatory molecules; its absence can lead to gut inflammation.^71–73^ Our findings align with the growing evidence that gut dysbiosis, characterized by deficits in protective microbes, can contribute to systemic inflammation, which may, in turn, affect the brain.^74–76^ The concurrent reduction of *Lachnospiraceae* (reflected by lower *Pseudobutyrivibrio*) and increase in *Bacteroides* and *Barnesiella* we observed fits the pattern of a dysbiotic microbiota in AD, as reported in prior studies.^77–79^ In fact, *Bacteroides* genus includes species that produce endotoxins and have been linked to neuroinflammation; its relative overgrowth may further exacerbate inflammatory tone.

Crucially, when we stratified by *APOE* genotype, we found that even *APOE* ε4 non-carriers (who have lower genetic AD risk) showed the core gut-amyloid relationship: higher Aβ linked to lower SCFA bacteria. This suggests that the gut microbiome’s association with amyloid is a general phenomenon not entirely dependent on *APOE*. However, in *APOE* ε4 carriers, the gut–Aβ associations were *more extensive*, with an even greater loss of beneficial microbes (e.g., *Clostridium*, *Barnesiella*, *Christensenellaceae* were significantly implicated only in ε4 carriers). Notably, we observed in ε4 carriers that *Barnesiella* and *Clostridium* were depleted with high Aβ, whereas in non-carriers *Barnesiella* wasn’t depleted. This aligns with a recent study by Tran *et al.,* which reported that *APOE* ε4 carriers (both humans and mice) had reduced levels of certain SCFA-producing taxa compared to non-carriers.^9^ Our data extend this by showing that *APOE* ε4 carriers with a heavy amyloid burden exhibit the most pronounced gut dysbiosis, suggesting that *APOE* ε4 and amyloid pathology may synergistically disrupt the microbiome. This supports the hypothesis that these SCFA-producing bacteria may contribute to the protective effect observed in *APOE* ε2 and *APOE* ε3 genotypes. In other words, *APOE* ε2/ε3 individuals might maintain a healthier microbiome (richer in taxa having anti-inflammatory properties), which could help limit amyloid or its consequences, whereas *APOE* ε4 individuals may lack this microbiome-mediated protection.

Our **mediation analysis** further strengthens the argument that the gut microbiome is not just correlated with, but potentially *causally involved* in, the *APOE*–amyloid link. We identified *Ruminococcus*, *Butyricicoccus*, *Clostridium*, and *Christensenellaceae* as significant mediators of the *APOE* ε4 effect on Aβ burden. These taxa are known for beneficial roles: *Ruminococcus*, *Butyricicoccus*, and *Clostridium* include fiber fermenters that produce SCFAs, and *Christensenellaceae* are associated with low inflammation and healthy metabolic profiles. The mediation suggests that *APOE* ε4’s association with higher brain amyloid burden is partly *through* its effect on lowering these microbes. To put it another way, if we could somehow restore *Ruminococcus*, *Butyricicoccus*, *Clostridium*, and *Christensenellaceae* in *APOE* ε4 carriers, we might attenuate some of *APOE* ε4’s pro-amyloid impact. This dovetails with literature highlighting the importance of protective microbes in maintaining the integrity of the blood-brain barrier (BBB) and regulating neuroinflammation.^80^ Anti-inflammatory bacteria, such as those producing SCFAs, can promote tight junction expression and reduce peripheral inflammation, thereby helping to protect the brain.^80^ Thus, the proposed mechanism linking *APOE* ε4 to amyloid accumulation likely involves SCFA-related pathways: *APOE* ε4 leads to a less favorable gut microbiome (perhaps due to *APOE* ε4-driven immunometabolic differences in the host gut environment), this microbiome produces fewer SCFAs, and the lack of SCFAs leads to a weakened BBB and more systemic inflammation, facilitating Aβ deposition and aggregation in the brain. Our findings provide empirical support for this mechanistic (often observed in mice^81,82^) model in humans.

The predictive functional analysis of the microbiome corroborated these ideas by showing a decrease in genes related to SCFA biosynthesis in high-amyloid individuals. Functions for producing acetate, butyrate, and propionate were all significantly down-represented. This adds a layer of evidence that it’s not just the names of bacteria that differ - the metabolic output of the microbiome is likely altered in a way that could impact the host. Lower abundance of key microbial enzymes like pyruvate dehydrogenase and propionyl-CoA synthetase in the gut suggests less SCFA is being made available. SCFAs have been shown to modulate microglial activation and astrocyte function in the brain; deficiency in SCFAs might tilt microglia towards a pro-inflammatory, less amyloid-clearing phenotype.^81^ Therefore, our results integrate well with the emerging paradigm that microbial metabolites are crucial intermediaries in the gut–brain axis influencing AD.^81^

From a broader perspective, our study highlights a few important points for the field of AD research and neurogastroenterology:

- **Host-microbiome interactions in AD:** We provide evidence that host genotype (*APOE*) and gut microbes interact to influence disease-related pathology. This underscores that AD risk is not purely intrinsic to the brain or genes; it may be modulated by potentially modifiable factors, such as the microbiome. *APOE* ε4 has long been seen as a “non-modifiable” risk factor, but if part of *APOE* ε4’s effect is microbiome-mediated, it opens the door to modifying the downstream consequences of *APOE* ε4 via diet, probiotics, or other microbiota-directed interventions.
- **Consistency with previous literature:** Our findings of reduced *Faecalibacterium* and *Ruminococcaceae* in association with AD biomarkers align with prior human studies, which have shown that these taxa are reduced in AD patients and in amyloid-positive cognitively impaired individuals.^77,83^ We also saw increased *Alistipes* and *Bacteroides* with amyloid, which has been reported (e.g., *Cattaneo et al.* found pro-inflammatory genera including some *Bacteroidetes* were higher with brain amyloid.^25^ The uniqueness here is linking it specifically to *APOE* groups and mediation.
- **Limitations:** We acknowledge several limitations. First, the cross-sectional design precludes definitive causal conclusions. While mediation analysis and biological plausibility suggest a causal chain, longitudinal studies are needed to confirm that microbiome changes precede and contribute to amyloid accumulation (there is some evidence from mouse models and one study in humans that gut changes can precede amyloid.^84^ Second, our microbiome data are based on 16S rRNA gene sequencing, which has limited taxonomic resolution and only infers function. Future studies should employ metagenomic shotgun sequencing and metabolomic profiling to directly measure microbial enzymes and metabolites (such as SCFAs and lipopolysaccharide) in *APOE*-stratified groups. Third, the cohort, while large for a PET study (n=227), is relatively healthy and mostly middle-aged; as such, results may differ in older, more clinically impaired populations. The generalizability to other ethnic groups or to those with advanced AD remains to be tested. Nonetheless, the FHS sample is community-dwelling and not enriched for AD, which is a strength in observing preclinical changes. Fourth, dietary factors and medication use were not extensively accounted for in this study; diet and medication have a strong influence on the microbiome and could confound the associations. However, one advantage is that all participants are from the same study site with similar regional diets, and we adjusted for BMI, which partially captures diet/lifestyle.

Despite these limitations, our study has several strengths, including the relatively large sample with both high-quality microbiome and PET imaging data, the use of robust statistical tools with multiple comparison correction, and replication of known microbiome-AD associations, which adds credibility to our novel findings about *APOE* interactions.

## Conclusion

This work provides compelling evidence of a link between *APOE* genotype, gut microbiome composition, and cerebral amyloid-β deposition. We discovered that *APOE* ε4-associated risk for AD may be partly mediated by perturbations in the gut microbiota, notably a deficiency in SCFA-producing bacteria that correlates with greater amyloid burden. These findings underscore a previously unappreciated connection between a major genetic risk factor for AD and modifiable microbial and metabolic pathways. The gut microbiome thus emerges as a key player in the complex network connecting *APOE* genotype to AD neuropathology.

Our results emphasize a possible role for gut-derived metabolites (like SCFAs) in protecting against amyloid-beta accumulation and suggest that *APOE* ε4 disrupts this protective link. This opens avenues for novel interventions: targeting gut bacteria that produce neuroprotective metabolites could potentially buffer the impact of *APOE* ε4 on the brain. Future studies should prioritize longitudinal designs and metagenomic analyses to validate these mediating relationships and to explore causality. If confirmed, manipulating the gut microbiome (through diet, probiotics, or fecal microbiota transplantation) may become a viable strategy for AD prevention, particularly in high-risk individuals. In conclusion, our study reveals a new facet of the gut-brain axis in Alzheimer’s disease, highlighting the gut microbiota as a potential therapeutic target at the intersection of genetics and neurodegeneration.

## Acknowledgements

The Authors thank Dr. Ramnik Xavier from the Broad Institute of MIT and Harvard, and the Center for Microbiome Informatics and Therapeutics (Massachusetts Institute of Technology, Cambridge, MA, USA) for providing access to the FHS microbiome data. We also thank Drs. Stanley Y Shaw, Charles De Carli, Keith Johnson, and Georges El Fakhri from the Broad Institute of MIT and Harvard.

## Data availability statement

All data supporting the findings of this study are publicly accessible through dbGap (Study ID: phs000007.v32.p13, https://www.ncbi.nlm.nih.gov/projects/gap/cgi-bin/study.cgi?study_id=phs000007.v32.p13).

## Authors contributions

B.F. conceived the study. J.J.H., C.L.S., S.S., R.S.V., and A.B. provided access to the data.

Y.N.W. performed all statistical analyses. Y.N.W. and B.F. wrote the manuscript. All authors discussed the results, provided feedback during the writing process, and commented on the final manuscript.

## Generative AI statement

The authors declare that no Generative AI was used in the creation of this manuscript.

